# PDBCleanV2: A Python Library for Generating Consistent Structure Datasets

**DOI:** 10.1101/2025.02.14.638326

**Authors:** Fátima Pardo-Avila, Liv Weiner, Paulina Cabral, Nicholas Corsepius, Frédéric Poitevin, Michael Levitt

## Abstract

The number of structures in the Protein Data Bank has grown rapidly in recent years. Nonetheless, comparing structures of a specific system often proves challenging due to variant nomenclature, origins from different species, presence of various ligands, and missing atoms or chains. To address these issues, we have developed PDBCleanV2, a Python package that enables users to create consistent datasets of structures, simplifying comparison among structures. This library generates individual files for each molecule in a structure file, corrects labeling errors, and standardizes chain names and numbering. Our aim is to provide researchers with consistent datasets that streamline their analysis. The source code, installation instructions, and tutorials can be found in PDBCleanV2’s GitHub repository and Zenodo accessible at https://github.com/fatipardo/PDBClean-0.0.2/ and https://doi.org/10.5281/zenodo.14014241, respectively.

## Introduction

Technological advances such as cryo-EM have led to a considerable increase in the quantity of macromolecular structures available in databases like the Protein Data Bank (RCSB PDB^1,2^). Even though typically only one structure exists for most unique polymer sequences in the RCSB PDB, many sequences have dozens of resolved structures (Supplementary Figure 1). A variable number of structures for a particular macromolecule could exist due to factors such as multiple organisms, differing conformational states, or diverse ligand interactions.

Comparing the available structures for a macromolecule can further our understanding of a system. However, this is often complicated by inconsistent nomenclature and structural variation, in structures which different research groups may have determined.

Specific macromolecule databases like GPCRs^3^, G proteins^4^, and the ribosome^5^ offer valuable resources. By accumulating all available information and structures, these databases present a more comprehensive understanding to researchers. However, creating such databases can be challenging for researchers interested in macromolecules not currently covered, given that it can be time-consuming and requires a bioinformatics background. To alleviate this problem, we have developed PDBCleanV2, which allows users to create consistent structure datasets in a user-friendly manner, even if they have limited bioinformatics experience.

Overall, PDBCleanV2 aims to ease the creation of consistent curated structure datasets, facilitating the discovery of novel insights from structure comparisons.

### Basics and Implementation

PDBCleanV2 is coded in Python and is open-source, available on GitHub (https://github.com/fatipardo/PDBClean-0.0.2/). The library requires BioPython^6^ and MUSCLE v5^7^; the complete list of dependencies is provided in the YML file. We recommend using Conda^8^ for PDBCleanV2 installation. The GitHub repository includes detailed installation instructions and tutorials (including ready-to-run Jupyter Notebooks^9^). The code has been developed and tested on MacOS.

### Workflow

A curated dataset comprises all available structures of a particular macromolecule of interest from the PDB. The curation process ensures that the same entities are labeled and numbered consistently across the dataset and rectifies any mislabeling errors from original structure submission to the PDB. Users can create a complete, curated dataset in five steps (Figure 1). Each step is described below. For additional instructions on using these functions, refer to our tutorial (https://bit.ly/PDBCleanV2).

**Figure 1.**
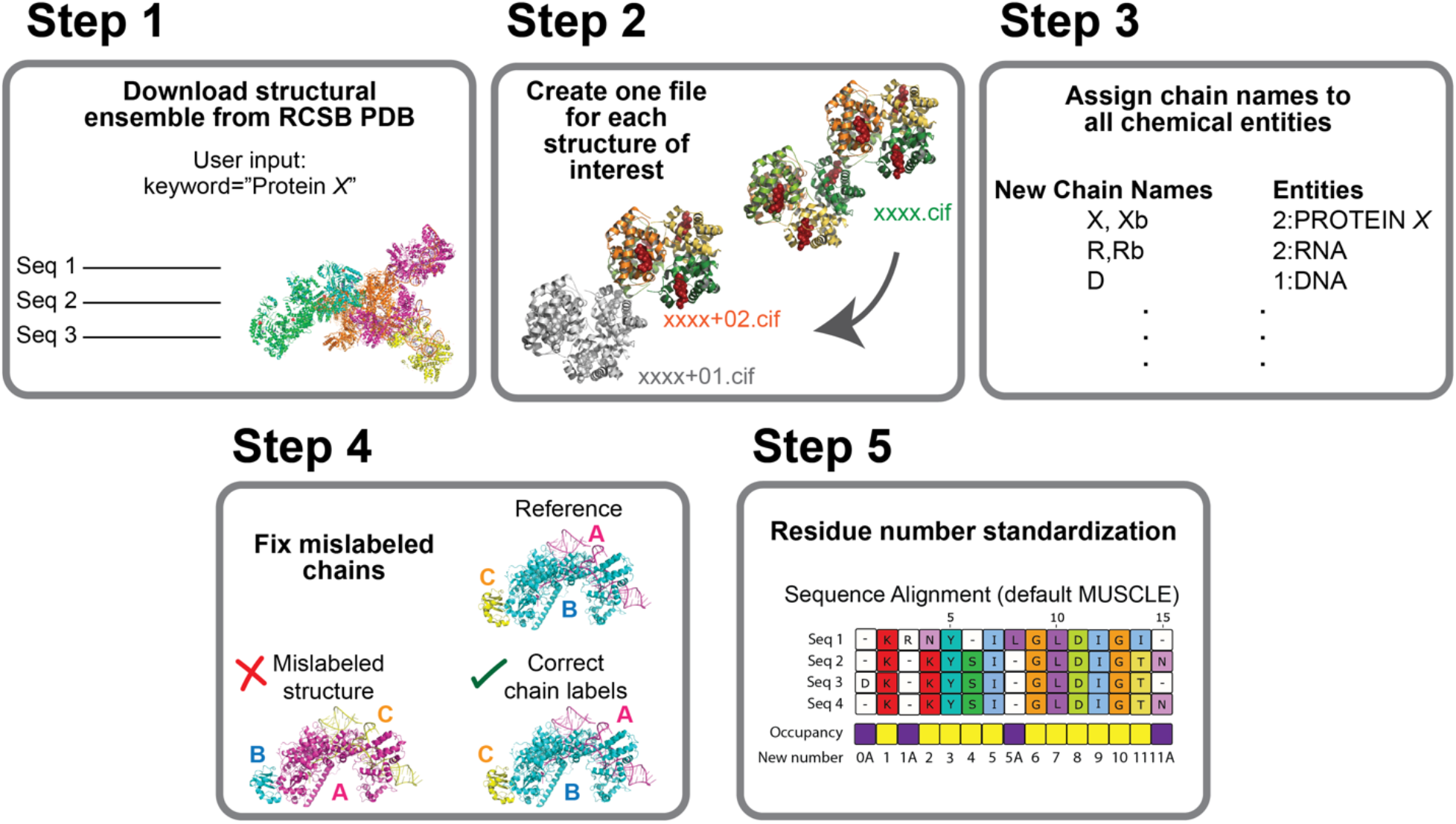
PDBCleanV2 workflow chart for creating a consistent, curated dataset of molecular structures.

### Step 1. Download Structures

PDBCleanV2 allows users to search for structures in the Protein Data Bank (PDB) using a reference sequence or keyword as a query. In the provided Jupyter Notebook in our tutorial (Step1.DownloadStructuralEnsembleFromRCSBPDB.ipynb), users need to input a keyword to download all relevant structures from the PDB. Initially, PDBCleanV2 downloads all available sequences from the PDB, then it searches for sequences that match the provided keyword. These matching sequences are subsequently used as queries to search the PDB for exact matches. Even incorrectly labeled structures are found in this step. PDBCleanV2 downloads all structures in CIF format (Figure 2).

**Figure 2.**
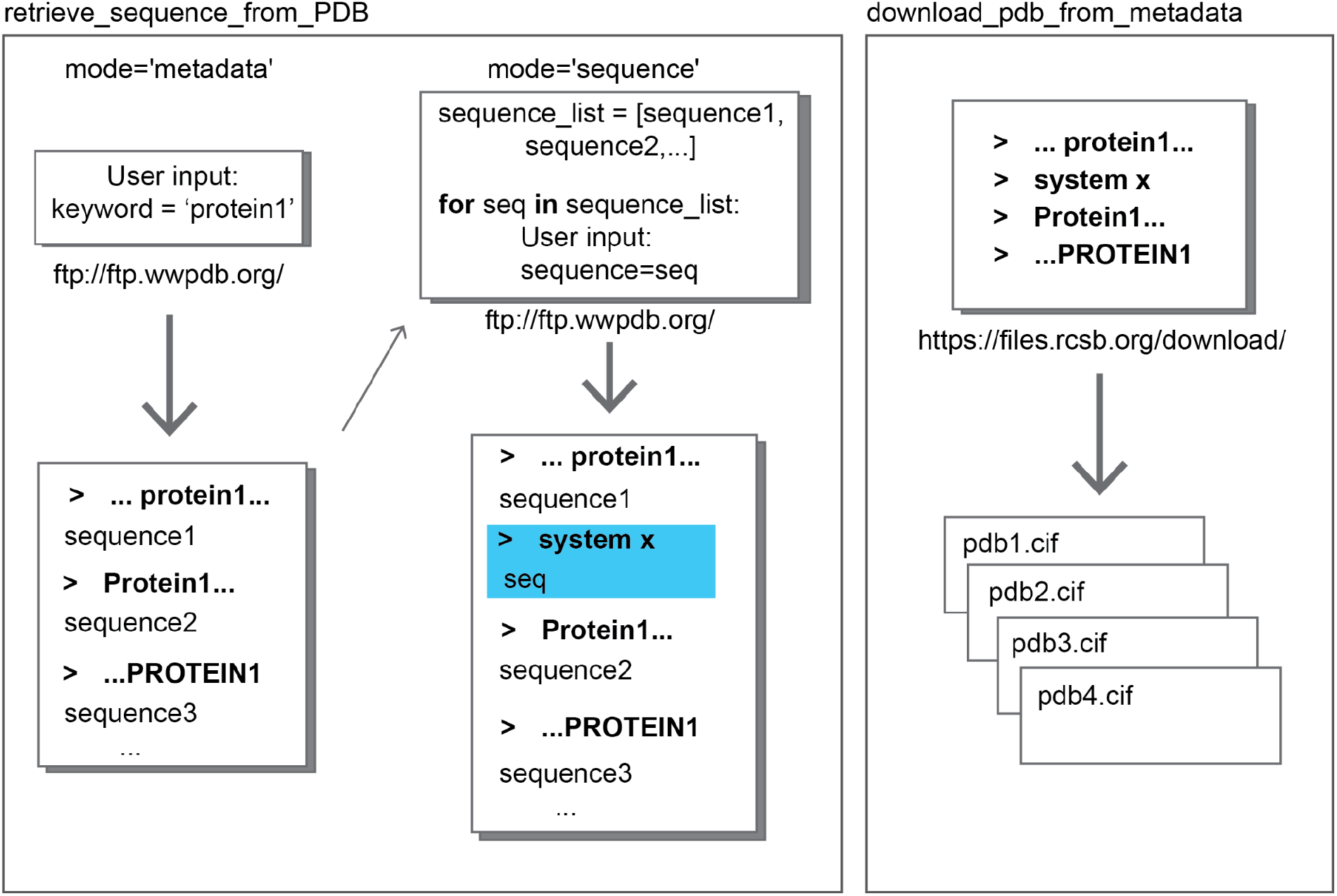
Step 1 - Downloading Structural Ensemble from RCSB PDB. Users can download a structural ensemble of any molecule of interest by inputting only a single keyword. PDBCleanV2 then searches for this keyword in the metadata (mode=“metadata”) of all the structures available in the PDB, retrieves all related sequences, and uses them as input for a subsequent search (mode=“sequence”). All the metadata of the retrieved sequences is input to the function “download_pdb_from_metadata,” which downloads the corresponding structures in CIF format from the RCSB PDB.

### Step 2. Generate a Single CIF File for Each Molecule of Interest

In a crystal structure, an asymmetric unit may contain multiple copies of the molecule under study. These copies are referred to as biological assemblies in the PDB file (SI Figure 2b). Each assembly’s atomic positions differ slightly, occupying unique positions within the asymmetric unit of the crystal^2^. Consequently, it is best to consider each molecule as a separate structure.

During the second step, PDBCleanV2 automatically separates all the molecules of interest within a single asymmetric unit, creating individual CIF files for each instance (Figure 3b). The number of data tags within these CIF files are also reduced and made consistent^10^. These tags contain critical structure information such as resolution, solution methods, authorship, among other things. Given that there are no guidelines specifying the tags to be included in a file, the data tags present may vary from structure to structure. PDBCleanV2 simplifies these files, preserving only the tags related to structure coordinates and authorship (Figure 3a).

**Figure 3.**
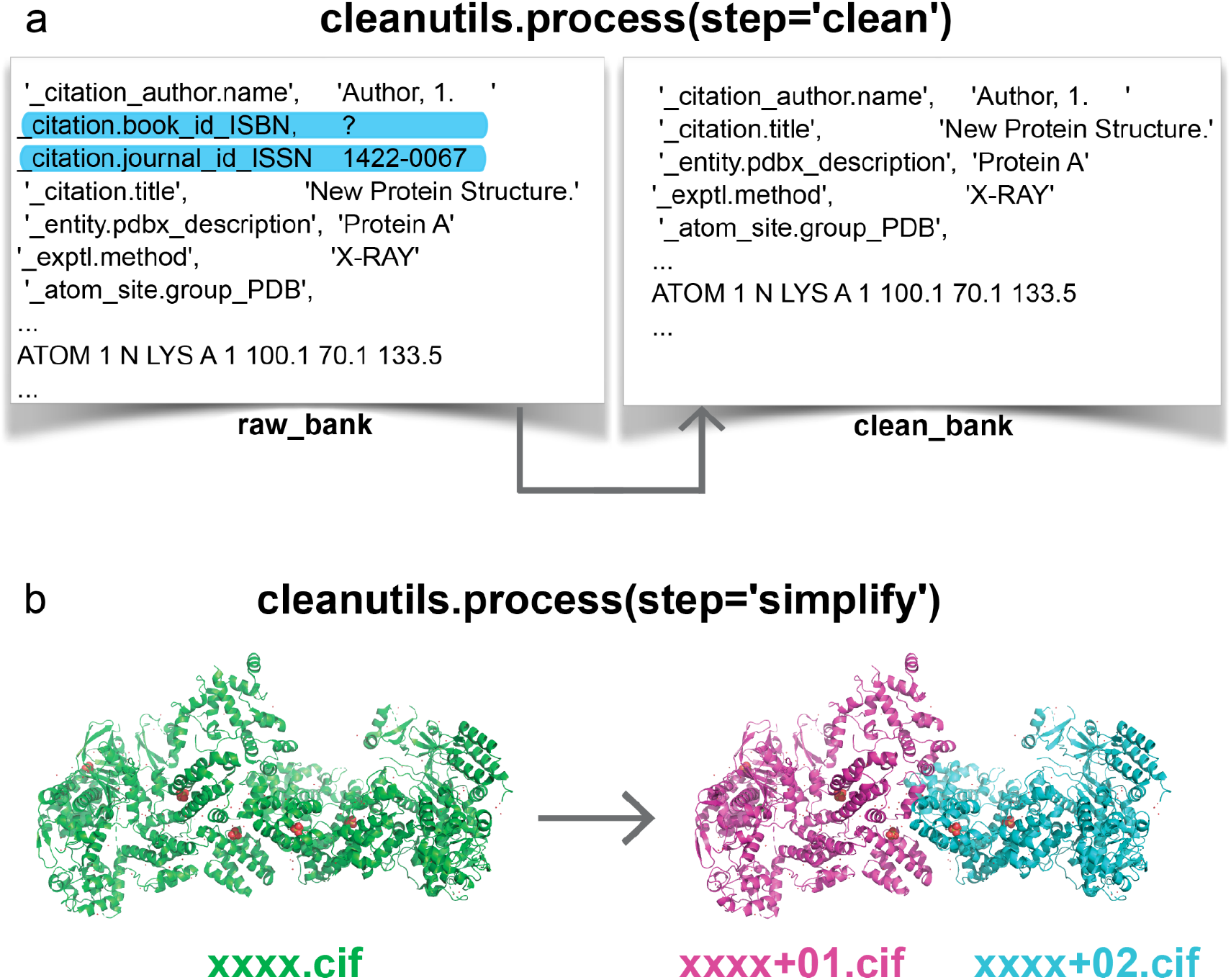
Step 2 - Cleaning CIFs and Creating One Structure Per Biological Assembly. a) The CIFs in the ensemble are standardized by removing unnecessary data labels from a CIF using the function “cleanutils.process(step=‘clean’)”. b) The function “cleanutils.process(step=‘simplify’)” separates all the molecules of interest into distinct CIFs.

### Step 3: Uniform Naming of Chains Across Structures

The PDBx/mmCIF format adopts the concept of entities, representing chemically distinct parts of the structure in the data file. A single entry may consist of multiple instances of a given entity, leading to several chains corresponding to the same entity. For example, ions could be distributed across multiple chains, or entities could contain copies of a protein chain. For example, a hemoglobin tetramer consists of two copies of an alpha and a beta chain (Supplementary Figure 2).

Step 3 involves creating a list of all entities in the dataset. This list also records the maximum number of chains corresponding to each entity across all structures in the dataset (Figure 4). Users are then required to assign unique names to the chains belonging to each entity. Although previous PDB formatting restricted authors to using a maximum of two characters when labeling chains, the CIF format imposes no such constraints. Hence, users now have the liberty to name chains more descriptively and intuitively. We recommend using concise yet descriptive chain names that also align with any existing naming conventions of the system under study. For instance, in the case of the ribosome, established nomenclatures already exist^11,12^, which could guide chain naming.

**Figure 4.**
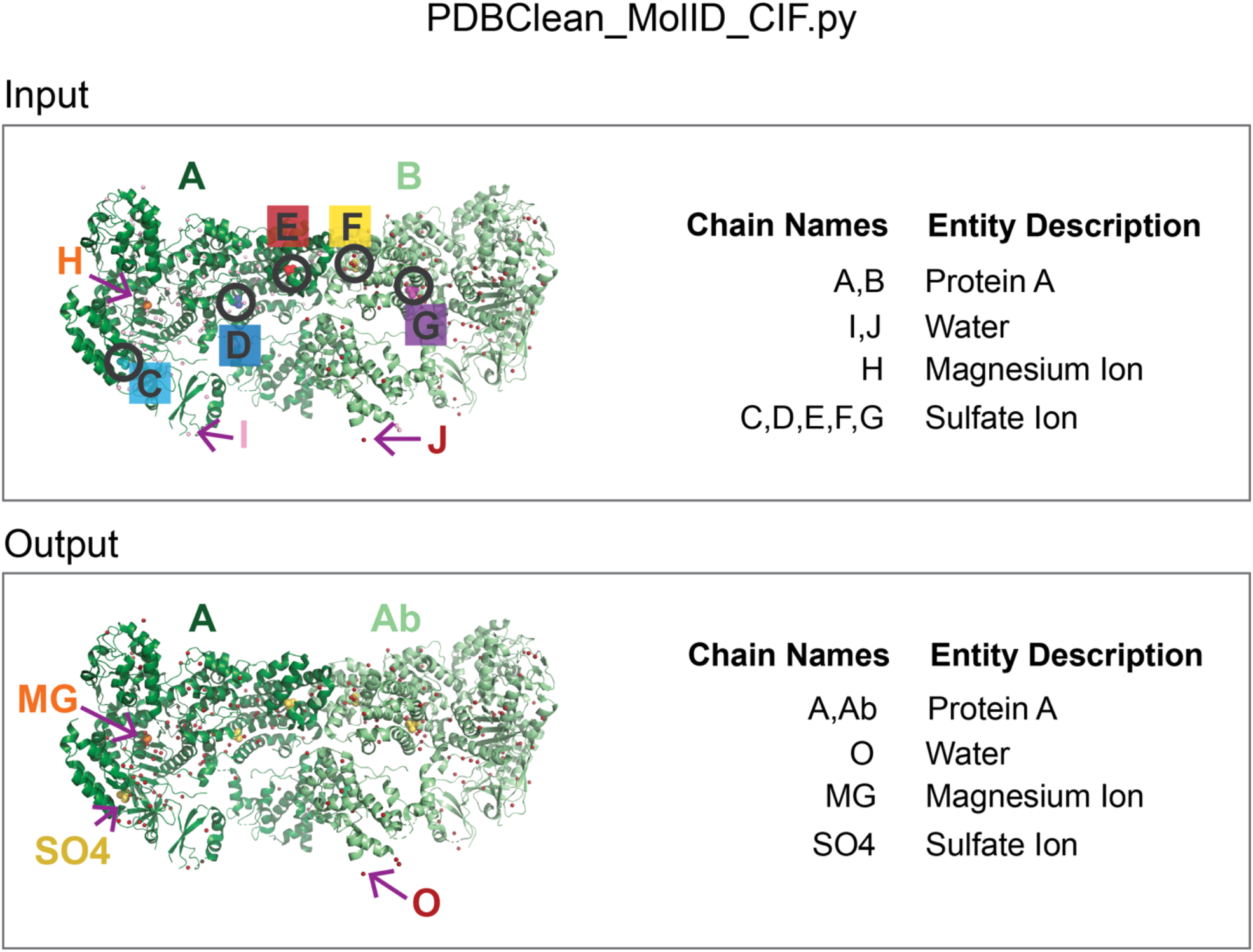
Step 3 - Name Assignment to All Entities in the Ensemble. Using PDBClean_MolID_CIF.py, users can generate a list of all entities in the structural ensemble and assign preferred names to these chains. The top panel shows an input structure and its corresponding entity list. The bottom panel shows a sample output, where the users have renamed the chain of each entity to make them more intuitive and easier to understand. Users might sometimes combine multiple chains, such as ions and water. In the case of ‘Protein A,’ two chains belong to this entity, meaning that there are two copies of ‘Protein A’ in at least one structure in the ensemble. In the case of ‘Sulfate Ion’, 5 separate sulfate ions previously in chains C, D, E, F, G are grouped in one entity. Concatenation of proteins is not recommended. Note that the new chain names are applied to all the structures in the ensemble.

In cases where multiple chains belong to the same entity, we recommend appending a suffix to the base name for each additional chain. For instance, for a chain ‘A’ with three copies, we suggest names ‘A’, ‘Ab’, and ‘Ac’. Lowercase suffixes prevent confusion when a chain ends with a number — a common occurrence in ribosome chain names.

Differentiating chain names is crucial, as PDBCleanV2 merges chains bearing the same name. However, this feature can prove beneficial for ions, as PDBCleanV2 can combine ions into a single chain.

PDBCleanV2 ensures a “self-consistent” dataset by uniformly labeling all structures in the set. When sharing the final structural dataset, it is essential that the key for chain names and entities also be provided.

Users should remember that entities can have multiple names. An efficient, automated procedure to group these entities and assign new chain names is currently unavailable. Inspecting the entities list helps identify outliers in the dataset. For instance, in the tutorial “Step3.1.AssignMolIDToEntitiesFoundInCIFfiles1.ipynb”, we demonstrated a dataset that includes an HLA Class II histocompatibility antigen bound to a fragment of the protein of interest, triosephosphate isomerase (TIM). Despite containing a short TIM fragment, the users might discard such structures from the final dataset since its dominant component is the HLA Class II histocompatibility antigen. While users ideally possess basic knowledge of the system they are curating to facilitate chain naming, experience with macromolecules/complexes can be gained during this step.

Although reviewing the entity list might be a time-consuming affair, it is crucial to establish a self-consistent dataset. It enables users to fathom the content of their curated dataset more deeply. The tutorials “Step3.1.AssignMolIDToEntitiesFoundInCIFfiles1.ipynb”, “Step3.2.AssignMolIDToEntitiesFoundInCIFfiles2.ipynb”, and “Step3.3AssignMolIDToEntitiesFoundInCIFfiles3.ipynb” elucidate working with the entities list and provide solutions to common problems that users might stumble upon during this curation step.

### Step 4: Chain Label Standardization

We’ve observed that some files may contain entities that are mislabeled. Therefore, it’s necessary to examine CIFs for any mislabeled entities. Our goal is to ensure that, within our curated datasets, all chains belonging to the same entity are consistently labeled across all structures. To achieve this, we compare the sequences of all chains in each structure to a set of reference sequences. Users can opt to provide these reference sequences, or PDBCleanV2 can automatically generate them.

A unique reference sequence is generated for each chain in the dataset. To establish a chain’s reference sequence, PDBCleanV2 collects all unique sequences assigned to that particular chain present in the dataset. It then counts the frequency of these sequences within the data set and selects the most prevalent one as the reference for that chain. This process is repeated for all chains.

Once the reference sequences have been obtained for each file in the dataset, PDBCleanV2 aligns the sequences of all chains in a structure against these reference sequences and scores the alignments. The score is determined by dividing the number of exact matches by the sequence length. These scores, along with the *hospital-resident matching algorithm*^13^, are used by PDBCleanV2 to decide the best chain name assignment (Figure 5).

**Figure 5.**
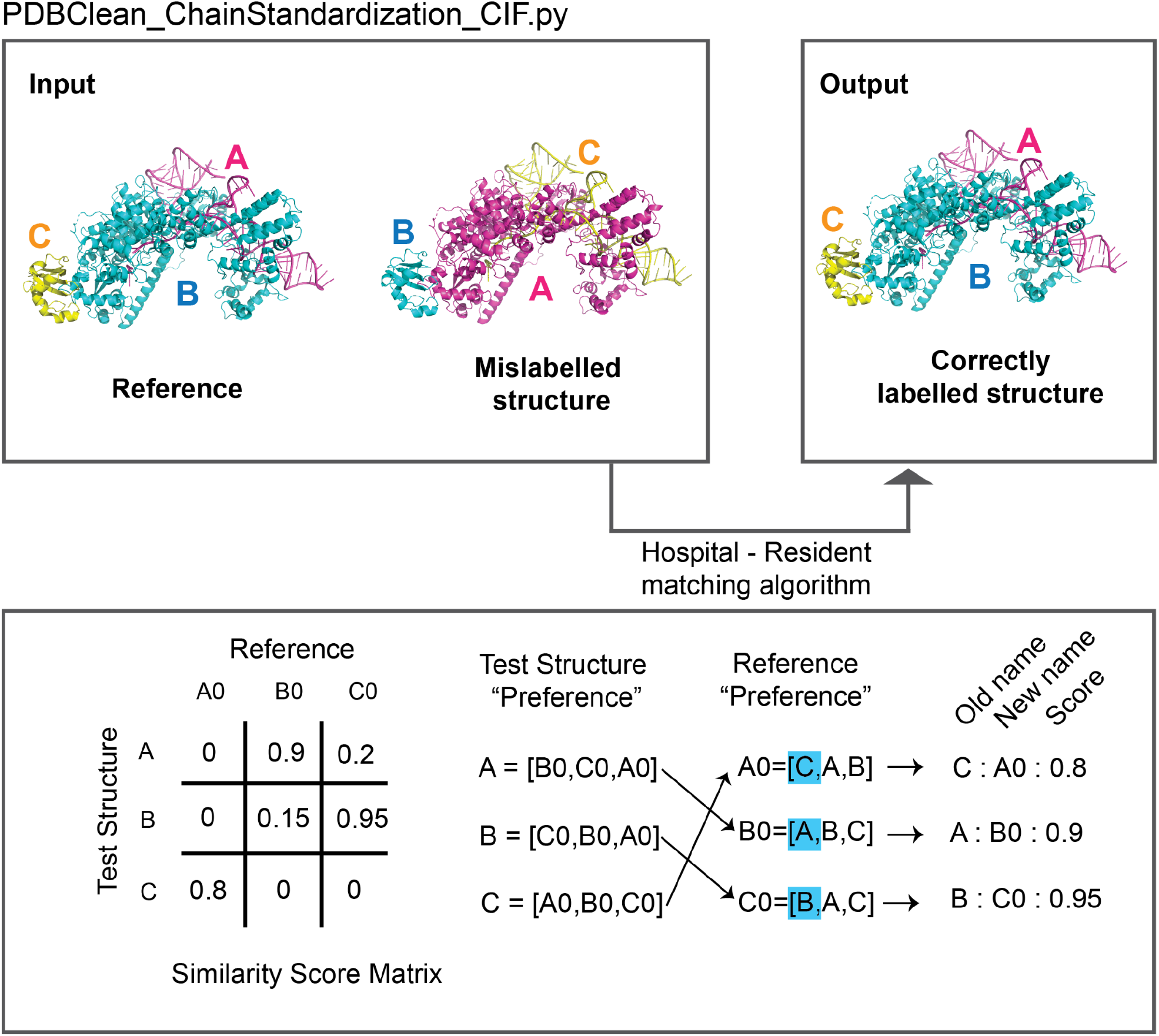
Step 4 - Chain ID Standardization. To correct any potential mislabelling, ‘PDBClean_ChainStandardization_CIF.py’ utilizes a sequence ensemble, a reference structure, and the *Hospital-Resident matching algorithm*^13^ to reassign accurate chain labels. The bottom panel demonstrates how the matching algorithm works. First, all the chains’ sequences in the test structure are compared against all the chains’ sequences of the reference, and similarity scores are calculated to fill a similarity score matrix. Then, for each chain in both the test and reference structures, a preference list is created by ranking based on the similarity scores. The preference lists are compared to obtain the matches used to rename the test structure. The output comprises the relabeled structures and a document containing the CIF ID, old and new chain names, and the similarity scores.

The output is a set of newly structured files with the new chain name assignments and a text file containing the CIF name, old and new chain assignments, and alignment scores. If the similarity score is low, users may wish to reject some changes. We’ve noted that unnecessary reassignments can occur when datasets contain structures from multiple organisms.

This process might require substantial CPU time for larger systems, as it involves aligning all chains in a macromolecular complex against every reference sequence. For instance, a system with 30 chains would need to perform roughly 30×30 pairwise alignments for each structure. Nevertheless, this process can potentially be parallelized. Users can split input files into multiple directories, run subsets of structures on PDBCleanV2, and apply the previously generated reference sequences. This procedure is detailed in the Jupyter Notebook tutorials “Step4.ChainIDStandardization.ipynb” and “Step4.2.ChainIDStandardization.ipynb”.

### Step 5. Standardization of Residue Numbering

Maintaining a consistent numbering system is crucial for comparing multiple structures. Nonetheless, it can be challenging due to reasons such as unresolved regions missing from a structure or a need to compare the same molecule from different organisms, which may have the same fold and significant residues in the same region but distinct sequences or residue insertions. PDBCleanV2 aims to assure that the same residue is consistently numbered across all the molecules in the curated dataset.

An alignment must be generated for each chain sharing the same name within the dataset. By default, PDBCleanV2 uses MUSCLE v.5 to generate the sequence alignments which are saved as fasta files. We advise users to verify the alignments prior to proceeding with the renumbering process. Users also have the option - and are encouraged - to provide their alignments, as in some instances, the use of a structural alignment or a combination of sequence and structural alignments may yield superior results.

A new numbering congruent for all structures is selected based on the multiple sequence alignment. To account for instances where only a few sequences exhibit insertions at certain positions, we only count positions where a residue is present in at least a certain percentage of the sequences in the dataset. The users can define what threshold suits their system the best. Figure 6 shows an example where we used 75% occupancy as the threshold. For positions that don’t meet this criterion, we append a letter to the number of the last counted residue (Figure 6).

**Figure 6.**
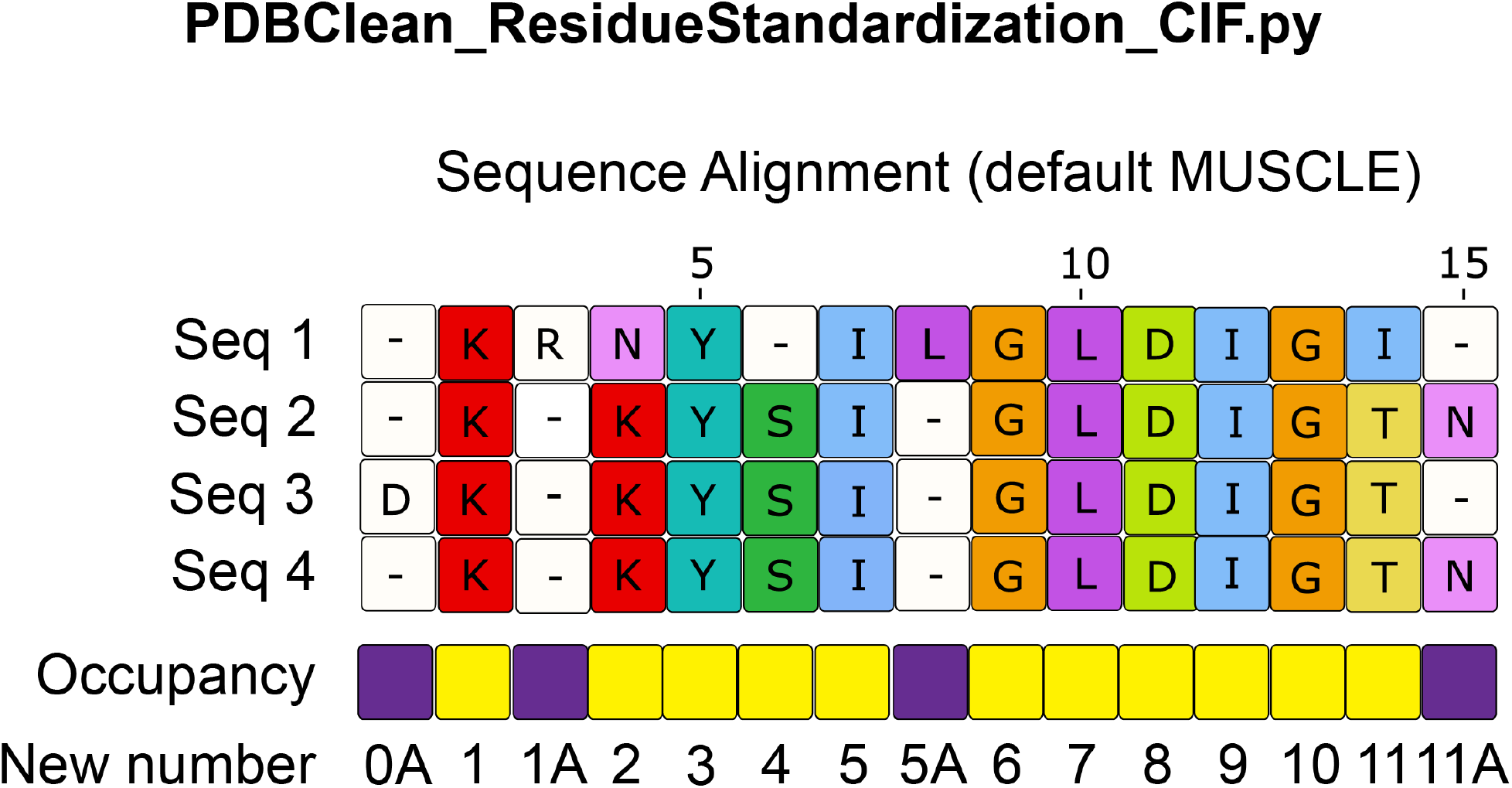
Step 5 - Residue ID Standardization. A multiple sequence alignment is conducted for each chain in the structural ensemble using MUSCLE v.5. Based on this alignment, a new numbering congruent for all structures in the dataset is chosen. For instances where only several sequences have insertions at certain positions, positions are only counted if a residue is present in at least a certain percentage of the sequences in the dataset. The user defines the occupancy threshold; we specified a 75% occupancy threshold in this example. Yellow squares show the positions that satisfy the threshold. If the condition isn’t met (i.e., residue missing in 75% of sequences, as marked by purple squares), a letter is added to the number of the last counted residue.

The output consists of a new set of structures with the revised residue numbers and a text file detailing the CIF name, chain names, and old and new residue numbers. The Jupyter Notebook tutorial “Step5.ResidueIDStandardization.ipynb” provides a demonstration of renumbering and the use of multiple sequence alignments in curated dataset creation.

### Tutorials and Curated Datasets

We prepared our tutorials as easy-to-use, ready-to-run Jupyter Notebooks. These serve to guide users through steps 1 to 5 as detailed in this manuscript and to demonstrate common challenges faced when generating a dataset. To ensure that these tutorials run smoothly and swiftly on personal computers within Jupyter Notebooks, they feature a small system: the enzyme triosephosphate isomerase (TIM). They illustrate how to execute certain functions using only a subset of TIM structures. The curated dataset is available to the public^14^, and we also provide the entity list and corresponding chain ID assignments.

## Conclusion

We’ve developed PDBCleanV2, a tool designed to assist researchers in creating self-consistent structure datasets. It provides a user-friendly interface and facilitates structure comparison. User involvement throughout the curation process enables datasets to be tailored to meet individual research needs.

The accompanying Jupyter Notebook tutorials demonstrate how to utilize our tool. Within these tutorials, we address users’ challenges in curating their datasets and provide potential solutions that optimize using PDBCleanV2 to its fullest extent. We also make available the curated datasets we have generated. They can be downloaded from our OSF profile^14^. Additionally, for each dataset, we generate a summary image that displays general features such as structure resolution distribution, publication year, structure-solving method, and origin organisms (generated using the “Analysis.SummaryPDBDataset.ipynb” notebook included in our GitHub repository).

We hope that users will leverage PDBCleanV2 as an initial stage in their research. It will allow them to learn from comparing all available structures of their selected biomolecules. Future PDBCleanV2 versions will include analysis tools, and we aim to provide the option to incorporate computer-generated models into the curated datasets.

## Key Points

- With PDBCleanV2, users can establish self-consistent structure datasets to compare structures easily.
- PDBCleanV2 generates individual files for each biological assembly in a structure file. It also corrects labeling errors and standardizes chain names and numbering.
- The source code is publicly accessible on GitHub. Detailed installation guidelines and Jupyter Notebook tutorials are provided.

## Supplementary Files

GitHub site with code and tutorials in Jupyter Notebooks: https://github.com/fatipardo/PDBClean-0.0.2/

OSF site with curated datasets: https://osf.io/9×8k2/

## Supplementary Figures

**Supplementary Figure 1.**
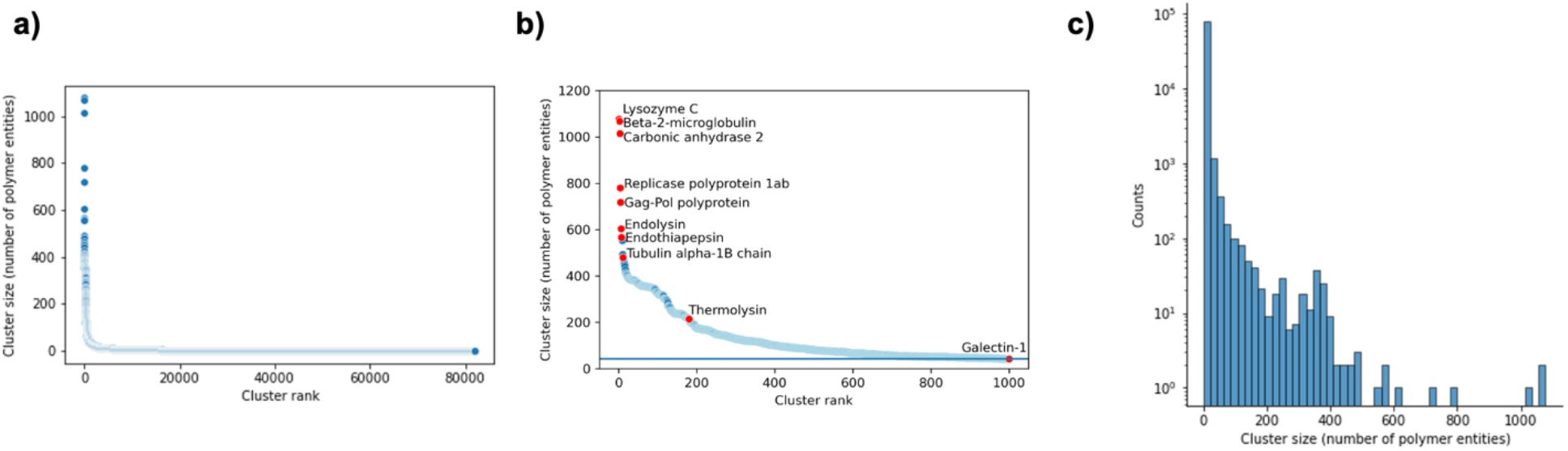
As of July 26th, 2023, the Protein Data Bank (PDB) contained 207,791 structures. Among these, approximately 90K were unique protein sequences, which were determined based on the number of protein clusters with 95% sequence identity. Part (a) displays the cluster size of all sequence clusters with 95% sequence identity, while part (b) shows that at least 1,000 sequence clusters contain 43 or more entries. Some clusters are highlighted in red and labelled with the associated protein name. Part (c) depicts a histogram of the unique polymer entities’ cluster size distribution, revealing that most unique sequence clusters contain fewer than ten entries.

**Supplementary Figure 2.**
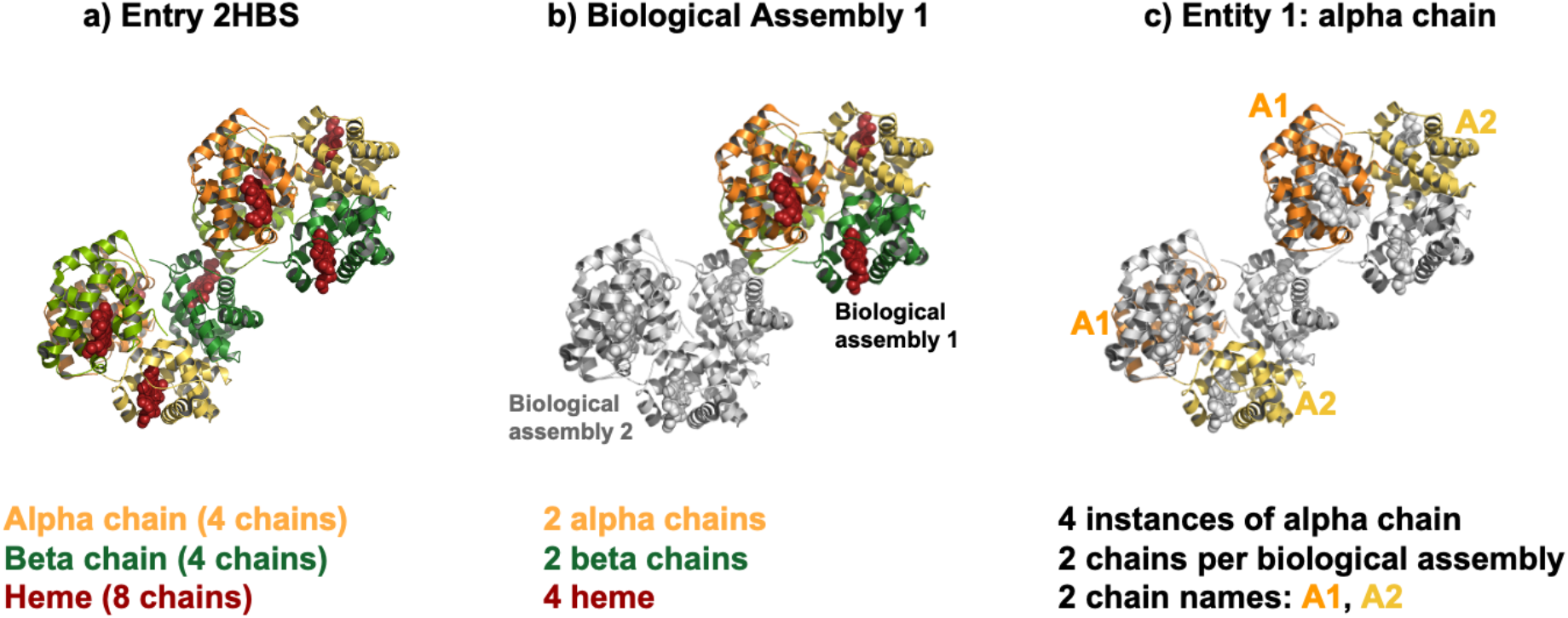
Understanding PDBCleanV2 Definitions. a) PDB entry 2HBS comprises two copies of sickle cell hemoglobin tetramers. These two copies of hemoglobin together form what is called the asymmetric unit. b) Each hemoglobin tetramer is categorized as a biological assembly. During step 2, PDBCleanV2 works to isolate each biological assembly, subsequently saving it as a distinct CIF. The diagrammatic representation of a biological assembly (colored) illustrates its components: two alpha chains (colored in orange and yellow), two beta chains (colored in forest green and lime green), and four separate heme molecules (colored firebrick red). c) Each component within the assembly is identified as an entity. For instance, Entity 1 is made up of the alpha chains, with each monomer containing two chains. PDBCleanV2 offers users the option to assign names to the chains. As there are two chains within each monomer, users are required to submit two names. If a single name is provided, PDBCleanV2 automatically concatenates the chains. This figure has been modified from the original^15^.

